# Rare genera differentiate urban green space soil bacterial communities in three cities across the world

**DOI:** 10.1101/2021.02.21.432167

**Authors:** Jacob G. Mills, Caitlin A. Selway, Laura S. Weyrich, Chris Skelly, Philip Weinstein, Torsten Thomas, Jennifer M. Young, Emma Marczylo, Sudesh Yadav, Vijay Yadav, Andrew J. Lowe, Martin F. Breed

**Affiliations:** School of Biological Sciences, The University of Adelaide, Adelaide, Australia; Department of Anthropology and Huck Institutes of the Life Sciences, Pennsylvania State University, USA; Healthy Urban Microbiome Initiative; Research & Intelligence, Public Health Dorset, Dorset County Council, UK; Environment Institute, The University of Adelaide, Adelaide, Australia; School of Public Health, The University of Adelaide, Adelaide, Australia; Centre for Marine Science and Innovation, School of Biological, Environmental and Earth Sciences, University of New South Wales, Sydney, Australia; College of Science and Engineering, Flinders University, Bedford Park, South Australia; Toxicology Department, Centre for Radiation, Chemical and Environmental Hazards, Public Health England, Chilton, Oxfordshire, UK; School of Environmental Sciences, Jawaharlal Nehru University, New Delhi, India

**Keywords:** microbial conservation, urban green space, vegetation complexity, soil bacteria, urban design

## Abstract

Vegetation complexity is potentially important for urban green space designs aimed at fostering microbial biodiversity to benefit human health. Exposure to urban microbial biodiversity may influence human health outcomes via immune training and regulation. In this context, improving human exposure to microbiota via biodiversity-centric urban green space designs is an underused opportunity. There is currently little knowledge on the association between vegetation complexity (i.e., diversity and structure) and soil microbiota of urban green spaces. Here, we investigated the association between vegetation complexity and soil bacteria in urban green spaces in Bournemouth, UK; Haikou, China; and the City of Playford, Australia by sequencing the 16S rRNA V4 gene region of soil samples and assessing bacterial diversity. We characterized these green spaces as having ‘low’ or ‘high’ vegetation complexity and explored whether these two broad categories contained similar bacterial community compositions and diversity around the world. Within cities, we observed significantly different alpha and beta diversities between vegetation complexities; however, these results varied between cities. Rare genera (< 1 % relative abundance individually, on average 35 % relative abundance when pooled) were most likely to be significantly different in sequence abundance between vegetation complexities and therefore explained much of the differences in microbial communities observed. Overall, general associations exist between soil bacterial communities and vegetation complexity, although these are not consistent between cities. Therefore, more in-depth work is required to be done locally to derive practical actions to assist the conservation and restoration of microbial communities in urban areas.

## Introduction

Microorganisms are important to every major biogeochemical process on Earth. They fix nitrogen, draw carbon-dioxide down from the atmosphere, weather rocks, decompose organic material, and, among many other things, form the base of the food web (Cockell & Jones 2009; Rousk & Bengtson 2014). Furthermore, microorganisms form symbiotic relationships with many plants and animals where they often have important roles in regulating host health (Rook et al. 2003; Rosado et al. 2018; Carthey et al. 2020). However, these ecosystem functions and services are being degraded by anthropogenic global change leading to climate, biodiversity and health crises. Urbanization in particular is linked to a public health crisis of rapidly rising non-communicable disease rates that are linked to losses of human exposure to microbial biodiversity (Rook et al. 2003; von Hertzen et al. 2011). Indeed, there have been repeated calls to conserve and restore microbial biodiversity (Blanco & Lal 2008; Bello et al. 2018; Mills et al. 2019; Carthey et al. 2020) due to the impact of human activities on ecosystem and human health (von Hertzen et al. 2011; Zuo et al. 2018).

One potential area where microbial communities could be conserved and restored is urban green spaces, and these areas are already used to help mitigate many issues that urbanization has on public health in general (Kabisch et al. 2017; Mills et al. 2019). Certain urban green space designs can reduce air pollution (Ferkol & Schraufnagel 2014; Xing & Brimblecombe 2020) and heat island effects (Aflaki et al. 2017), while potentially restoring microbial biodiversity to benefit ecosystem services (Hoch et al. 2019; Joyner et al. 2019). Indeed, restoring the urban microbiota by planting native vegetation could improve the exposure to microbes that humans need for immune training and regulation, thus contributing to reducing the immune disease prevalence found in cities (Rook et al. 2003; von Hertzen et al. 2011; Mills et al. 2017). Further, there is growing evidence that environmental microbiota can transfer readily to humans through inoculated play-ground media (Hui et al. 2019) or by simply using green spaces (Selway et al. 2020), and that vegetation type or diversity near the home is associated with human microbial diversity (Pearson et al. 2020).

Community characteristics of vegetation, such as species richness and functional diversity, are closely linked to microbial communities, including urban soils (Hui et al. 2017; Laforest-Lapointe et al. 2017; Mills et al. 2020). Soil in revegetated urban areas have microbial communities more representative of remnant areas compared to typical Victorian-era green spaces, such as lawns (Baruch et al. 2020; Mills et al. 2020). These associations are likely driven by plant-microbe-soil chemistry feedback loops (Berendsen et al. 2012; Fierer 2017). However, this evidence for the relationship between vegetation complexity of urban green space and their associated soil microbiota remains limited. As such, here we build on our earlier work in a single city (Mills et al. 2020) to focus on the association of vegetation complexity and soil bacterial communities both within and between three cities across different regions of the world.

## Methods

### Study sites

We focused our study on urban green spaces that represented ‘low’ or ‘high’ complexity vegetation in the cities of Bournemouth, UK; Haikou, China; and the City of Playford (hereafter known as Playford), Australia (Figure 1). Bournemouth has a ‘marine’ climate with short dry summers and heavy precipitation during mild winters, Haikou has a ‘humid subtropical’ climate, and Playford has a ‘Mediterranean’ climate. The green spaces were categorized as ‘high’ (i.e., remnant woodlands, revegetated woodlands, or regenerated woodlands) or ‘low’ (i.e., lawns, vacant lots, or parklands) complexity vegetation based on the diversity and structure of their vegetation (Figure 1). These two categories were based on our previous quantification of vegetation diversity (i.e., plant species richness) and structure (i.e., layers of plant growth-forms creating 3D structure) in the urban green space sites of Playford (quantified in Mills et al. 2020). In each city, we selected six sites of ‘low’ and six sites of ‘high’ complexity vegetation (example photos in Figure 1) by using local knowledge of existing urban green space vegetation types. Within each site, a 25 × 25 m quadrat in a NSEW orientation was sampled, with geo-references and photos taken at the SW corner.

**Figure 1.**
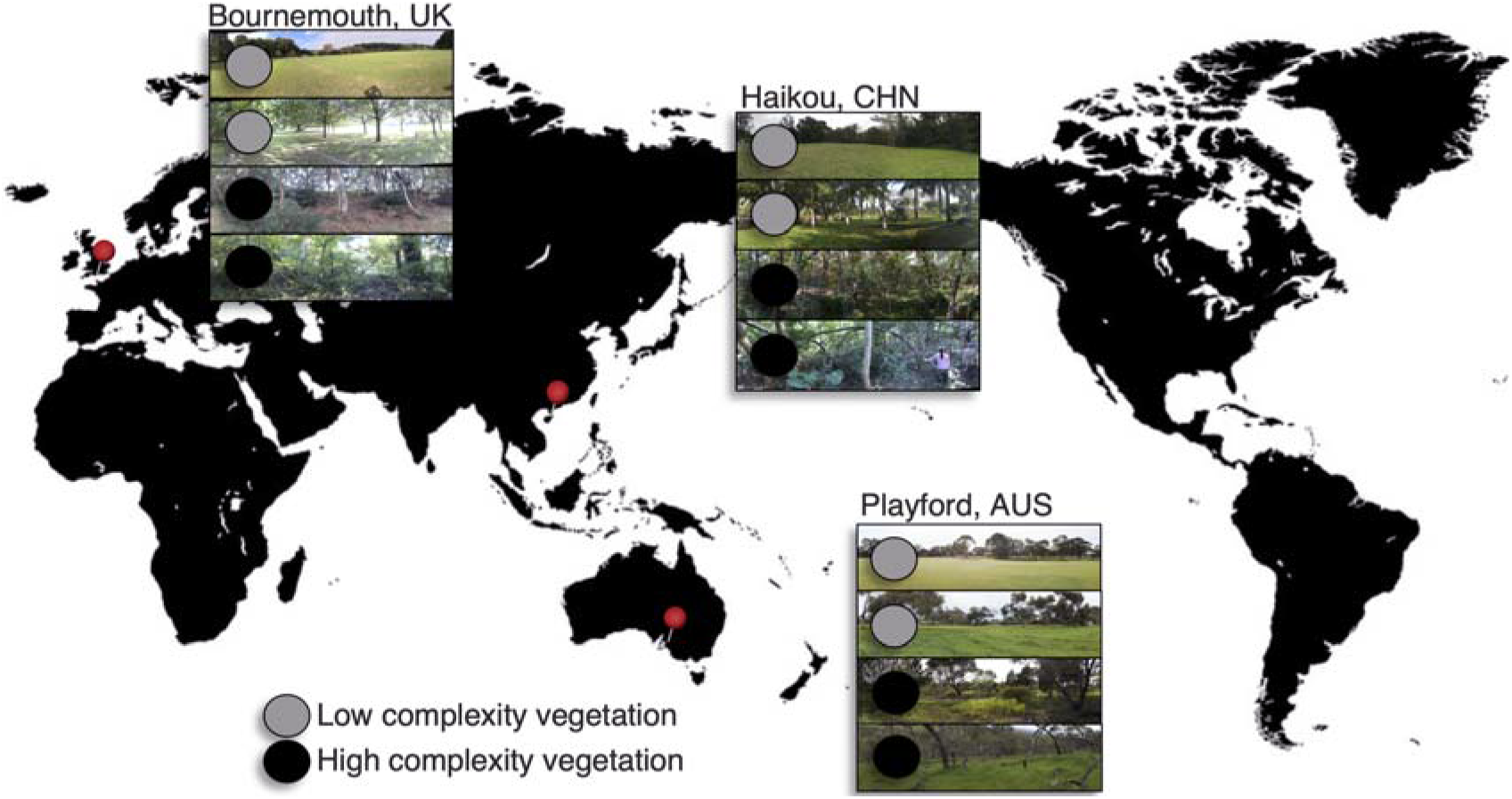
‘Low’ and ‘high’ complexity vegetation urban green spaces were sampled in Bournemouth, UK; Haikou, CHN; and Playford, AUS.

### Soil sampling

Soils were sampled for DNA extraction according to the Biomes of Australian Soil Environments (BASE) project protocol (Bissett et al. 2016) in September and October 2016. In brief, 100-200 g of soil from nine points within the quadrat were randomly sampled, pooled, and homogenized. From this pooled sample, 50 g were stored at −20°C until microbial analysis. The Bournemouth and Haikou samples have not been analyzed previously. The Playford samples are a subset of those reported in (Mills et al. 2020), including all samples except those from Parkland sites.

### Microbial community analysis

Soil DNA was extracted (one extraction from 0.2 g of each 50 g sample) using the DNeasy Powerlyser soil kit (QIAGEN) in the country of sampling, as per the manufacturer’s instructions. Extraction blank controls were not used; however, high biomass samples, such as soil, are less susceptible to contamination compared to those of low biomass and are therefore unlikely to produce results heavily swayed by contaminants (Velásquez-Mejía et al. 2018; Eisenhofer et al. 2019). Extracted DNA was then shipped to the University of Adelaide for downstream analysis as per Selway et al. (2020). Briefly, the bacterial 16S rRNA V4 gene region was amplified using primers 515F and barcoded 806R (Caporaso et al. 2011; Caporaso et al. 2012), and PCR components and cycling conditions were followed as previously described (Selway et al. (2020). PCR products were pooled into groups of approx. 30 samples at equimolar concentrations. Pools were cleaned (AxyPrep Mag Clean-up kit; Axygen Scientific), quantified and pooled together into a final sequencing pool before sequencing the DNA at the Australian Genome Research Facility using a 2 × 150 bp kit on an Illumina MiSeq.

In QIIME2 (v 2018.8), DNA sequences were merged, trimmed to 150 bp, and quality filtered (>Q4), and resulting sequences were denoised with deblur (Amir et al. 2017) to create amplicon sequence variants (ASVs), as previously described (Selway et al. (2020). Representative ASVs were assigned to the SILVA database (version 132). To remove laboratory contaminant sequences, ASVs were identified from PCR negative controls using the prevalence method within the *decontam* package (v 1.8.0; Davis et al. 2018) in R (v 4.0.0; RCoreTeam 2019) and with a threshold probability of 0.5. Any identified contaminants were removed from all biological samples before downstream analysis. Additionally, ASVs assigned to mitochondria, chloroplast, Archaea, or ‘unknown’ kingdom were also removed, and ASVs with fewer than ten reads across all samples in the dataset were excluded. Post-filtering, there were at least five ‘low’ and five ‘high’ complexity vegetation replicates for each city (see sample metadata via links in ‘*Data access*’).

### Statistical analyses

All statistics were done in R (v 4.0.0; RCoreTeam 2019). ASVs were agglomerated to genus level for statistical analysis using the ‘tax_glom’ function of the *phyloseq* package (v 1.32.1; McMurdie & Holmes 2013). During the genus agglomeration, all unresolved taxa at genus level, i.e. ‘NA’ or ‘blank’, were removed.

Before alpha diversity was calculated, the agglomerated genus level data was rarefied to 2,396 reads with the ‘rarefy_even_depth’ function of the *phyloseq* package. Alpha diversity was calculated as observed genus richness and Shannon’s diversity with the ‘estimate_richness’ function in *phyloseq* and Faith’s phylogenetic diversity was calculated with the ‘pd’ function of the *picante* package (v 1.8.1; Kembel et al. 2010). We used generalized linear mixed models (GLMMs) to test for difference in alpha diversity by crossing the fixed factors of ‘city’ and ‘vegetation complexity’ and nesting the random factor of ‘site’ within ‘city’. GLMMs were done with the ‘glmer’ function of the *lme4* package v 1.1-25 (v 1.1-25; Bates et al. 2007). Distributions for the GLMMs were Poisson for observed genus richness (count data) and Gamma for Faith’s phylogenetic diversity and Shannon’s diversity (positive, non-integer, non-parametric data). The Poisson GLMM was tested for over-dispersion (result: ratio = 0.46). Main effects of the GLMMs were tested by Type II Wald Chi^2^ tests with the ‘Anova’ function of the *car* package (v 3.0-10; Fox et al. 2012). Pairwise comparisons of ‘city’ and ‘vegetation complexity’ combinations were tested by z-tests with Holm-Bonferroni P-adjustment with the ‘glht’ function of the *multcomp* package v 1.4-15 (v 1.4-15; Hothorn et al. 2014).

Ordinations of beta diversity were done with the ‘ordinate’ function in *phyloseq*. Ordinations were based on unrarefied data in principal coordinates analysis (PCoA) with Bray-Curtis and Jaccard distance matrices. We used PERMANOVA, with 999 iterations, with the ‘adonis’ function of the *vegan* package (v 2.5-6; Oksanen et al. 2017) to test the model of ‘vegetation complexity’ nested within ‘city’. Pairwise comparisons between nested vegetation complexities (e.g. Bournemouth Low vs. Bournemouth High) were tested by PERMANOVA with 999 iterations with the ‘pairwise.adonis2’ function of the *pairwise.adonis* package (v 0.0.1; Arbizu 2017).

We created a relative abundance stack plot by converting the rarefied genus abundances to percentages. All genera with total rarefied sequences across all samples being less than 1 % of total rarefied sequences were pooled into a single group named ‘< 1 % abund.’. The less than 1 % cut-off was determined by a rank-abundance curve of percentage abundance across all samples (Figure S1). We tested for differentially abundant bacterial genera between ‘low’ and ‘high’ complexity vegetation sites within each city. Log-2 fold-change measurement of bacterial genera was done using the ‘DESeq’ function of the *DESeq2* package in *phyloseq* (v 1.28.1; Love et al. 2014). *DESeq2* does an internal normalization, where the count of each genus within a sample is divided by the mean of that genus across samples. Differentially abundant genera (alpha = 0.05) were plotted into heatmaps using the ‘pheatmap’ function of the *pheatmap* package (v 1.0.12; Kolde & Kolde 2015). The differential abundance heatmap scale represents the mean abundance of each genus across samples as 0 with ± 3 standard deviations. The differential abundance heatmap rows and columns were clustered based on Manhattan distance to most efficiently arrange the grid. The heatmap trees represent how closely related a row or column are, not taxa, based on the scale in each cell. Unclassified genera were not included in the heatmap.

## Results and Discussion

### Each city had distinct soil microbial communities

We compared soil bacterial genera between cities and found that the communities were quite distinct from each other, regardless of vegetation complexity, both in terms of alpha diversity (observed genus richness, Chi^2^ = 28.67, Pr(>Chi^2^) < 0.001; Faith’s phylogenetic diversity, Chi^2^ = 21.02, Pr(>Chi^2^) < 0.001; Shannon’s diversity, Chi^2^ = 21.80, Pr(>Chi^2^) < 0.001; Figure 2a) and beta diversity distances (Bray-Curtis, F = 19.61, Pr(>F) = 0.001; Jaccard, F = 10.68, Pr(>F) = 0.001; Figure 2b). Further, beta diversity at the ASV-level had similar patterns to the genus-level results; however, the data were over-dispersed (i.e., significantly more variable than predicted for the model) and therefore not used further (Figure S2). These differences between cities were expected given their differences in geography and climate, where for example, temperature, aridity, and distance from the equator vary and each are strong predictors of soil microbial diversity (DelgadoLBaquerizo et al. 2018). Moreover, strong biogeographic zoning and distance-decay relationships have previously been observed for urban soil bacterial communities across ten cities within China (Yang et al. 2021).

**Figure 2.**
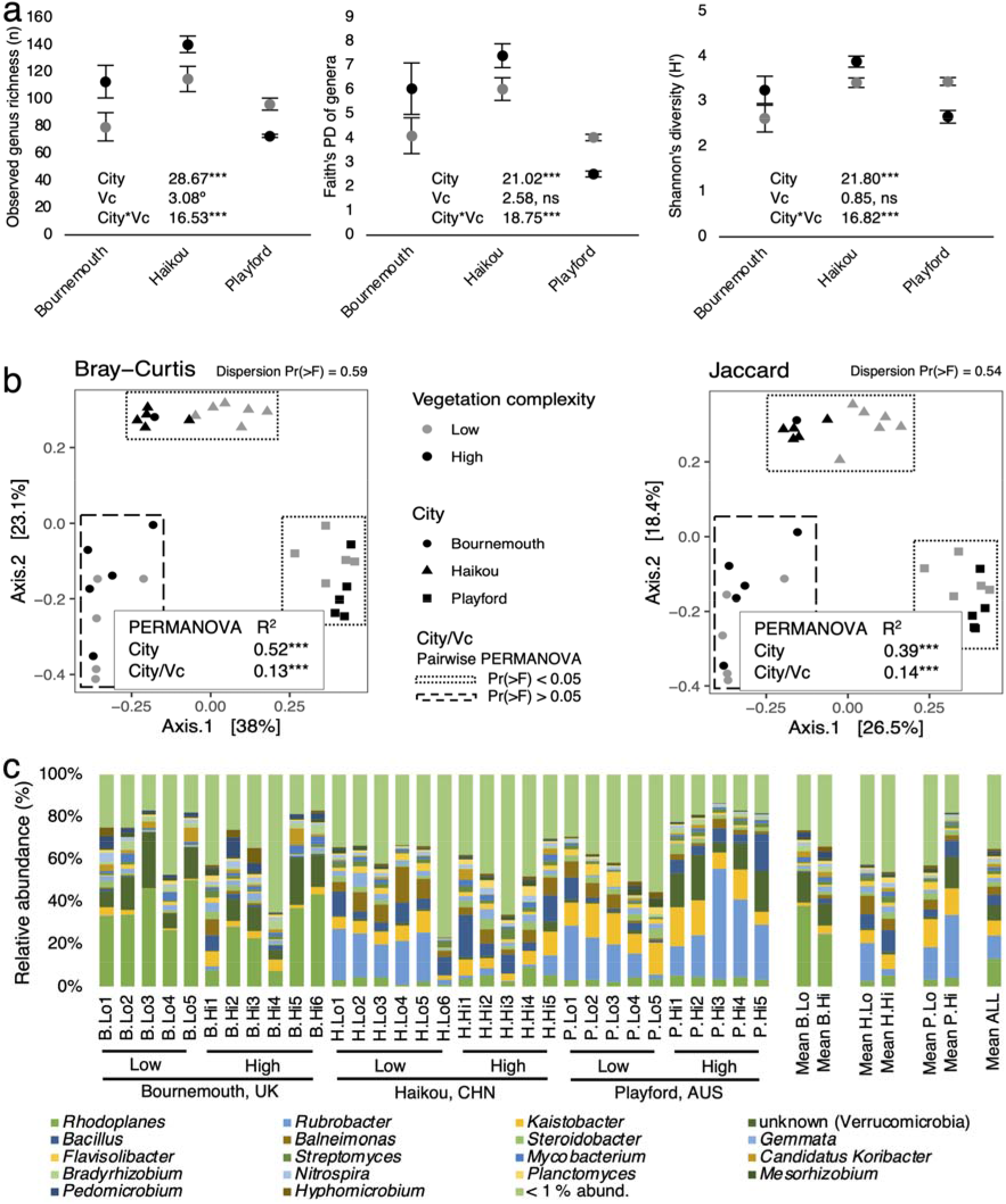
**(a)** ‘City’ by ‘Vegetation complexity’ (Vc) for the alpha diversity GLMMs on observed genus richness, Faith’s phylogenetic diversity (PD), and Shannon’s diversity. Results are Chi^2^ values from Type II Wald Chi^2^ tests on the GLMMs followed by significance codes for Pr(>Chi^2^). See Table 1 for pairwise results. Significance codes: ‘ns’ not significant; ‘°’ P < 0.10; ‘*’ P < 0.05; ‘**’ P < 0.01; ‘***’ P < 0.001. **(b)** PCoAs of soil bacterial genus communities in urban green spaces by Bray-Curtis and Jaccard distance. Main PERMANOVA test with 999 iterations of ‘Vegetation complexity’ (Vc) nested within ‘City’ (distance ~ City/Vc); R^2^ and P-value significance codes (‘***’, Pr(>F) < 0.001). Within city ‘Vegetation complexity’ differences were tested with pairwise PERMANOVA. Cities surrounded by dotted boxes were significantly different between their ‘low’ and ‘high’ vegetation complexity green spaces. Cities surrounded by dashed boxes were not significantly different between their ‘low’ or ‘high’ vegetation complexity green spaces. For detailed main and pairwise PERMANOVA results see Table 2. **(c)** Relative abundance (%) of soil bacterial genera across all sites. Genera read left to right by rows in the legend and correspond to bottom to top in the stack plot.

### ‘Low’ and ‘high’ complexity vegetation soils have similar diversity

We next compared diversity of sites with ‘low’ versus ‘high’ vegetation complexity within all three cities. In Bournemouth, ‘high’ complexity vegetation green spaces were significantly more diverse than ‘low’ complexity spaces for their bacterial genera (observed genus richness, z = 3.17, P = 0.014; Faith’s PD of genera, z = −2.93, P = 0.034), (Table 1 & Figure 2a). In contrast, alpha diversity of bacterial genera in soil from Playford and Haikou for all three measures were non-significantly different between the ‘low’ and ‘high’ complexity vegetation (Table 1). However, there was a significant interaction between ‘city’ and ‘vegetation complexity’ for all three alpha diversity measures (Figure 2a). This interaction was caused by the Playford soils being lower in diversity in the ‘high’ complexity vegetation soils relative to the ‘low’ complexity soils, whereas diversity was higher in these ‘high’ sites in Bournemouth and Haikou.

**Table 1.**
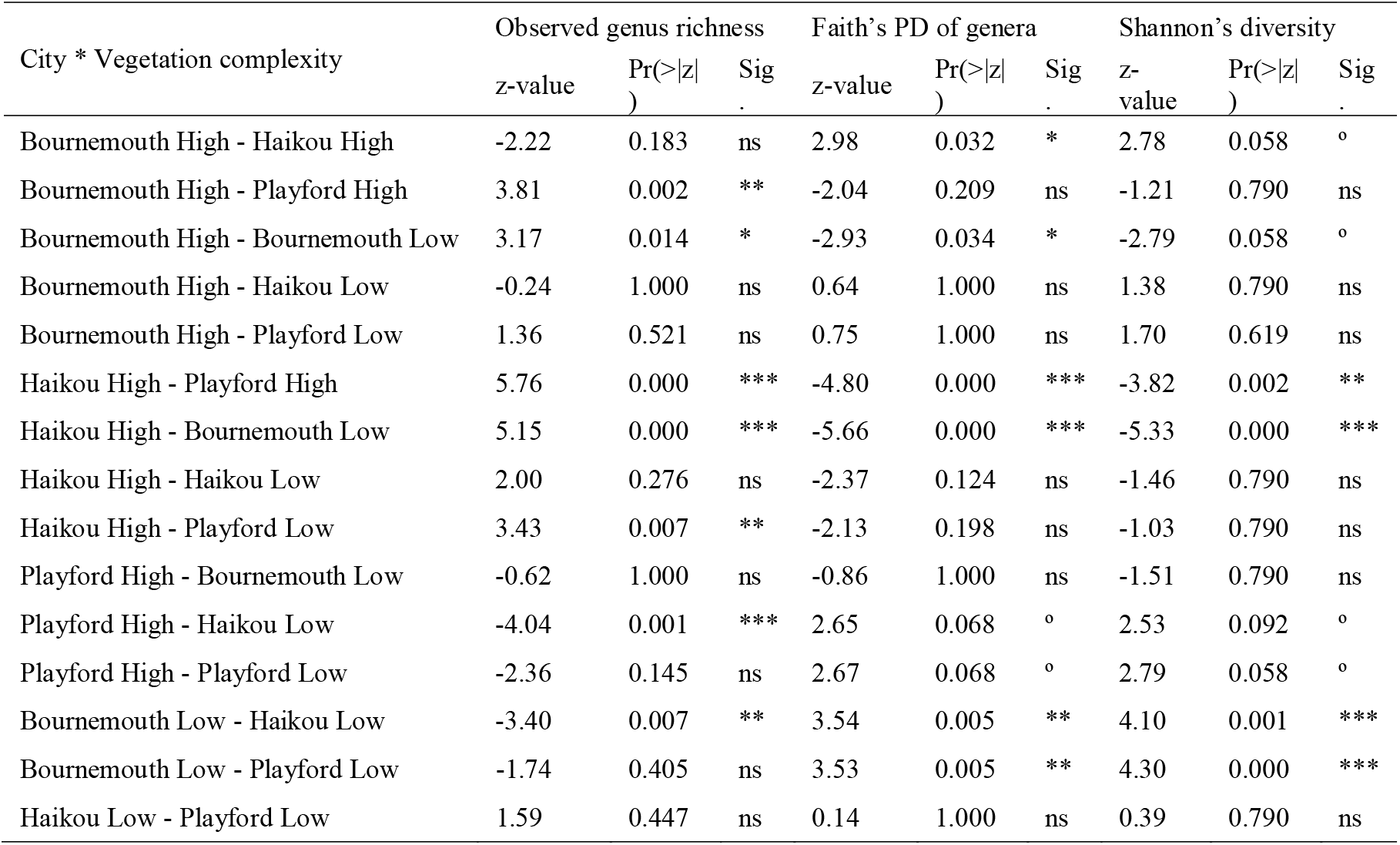
Pairwise alpha diversity – observed genus richness, Faith’s phylogenetic diversity (PD), and Shannon’s diversity – of soil bacterial genera under the GLMM interaction of ‘City’ by ‘Vegetation complexity’. Significance codes: ‘ns’ not significant; ‘°’ P < 0.10; ‘*’ P < 0.05; ‘**’ P < 0.01; ‘***’ P < 0.001.

| City * Vegetation complexity | Observed genus richness |  |  | Faith’s PD of genera |  |  | Shannon’s diversity |  |  |
| --- | --- | --- | --- | --- | --- | --- | --- | --- | --- |
|  | z-value | Pr(>\|z\|) | Sig. | z-value | Pr(>\|z\|) | Sig. | z-value | Pr(>\|z\|) | Sig. |
| Bournemouth High - Haikou High | -2.22 | 0.183 | ns | 2.98 | 0.032 | * | 2.78 | 0.058 | ° |
| Bournemouth High - Playford High | 3.81 | 0.002 | ** | -2.04 | 0.209 | ns | -1.21 | 0.790 | ns |
| Bournemouth High - Bournemouth Low | 3.17 | 0.014 | * | -2.93 | 0.034 | * | -2.79 | 0.058 | ° |
| Bournemouth High - Haikou Low | -0.24 | 1.000 | ns | 0.64 | 1.000 | ns | 1.38 | 0.790 | ns |
| Bournemouth High - Playford Low | 1.36 | 0.521 | ns | 0.75 | 1.000 | ns | 1.70 | 0.619 | ns |
| Haikou High - Playford High | 5.76 | 0.000 | *** | -4.80 | 0.000 | *** | -3.82 | 0.002 | ** |
| Haikou High - Bournemouth Low | 5.15 | 0.000 | *** | -5.66 | 0.000 | *** | -5.33 | 0.000 | *** |
| Haikou High - Haikou Low | 2.00 | 0.276 | ns | -2.37 | 0.124 | ns | -1.46 | 0.790 | ns |
| Haikou High - Playford Low | 3.43 | 0.007 | ** | -2.13 | 0.198 | ns | -1.03 | 0.790 | ns |
| Playford High - Bournemouth Low | -0.62 | 1.000 | ns | -0.86 | 1.000 | ns | -1.51 | 0.790 | ns |
| Playford High - Haikou Low | -4.04 | 0.001 | *** | 2.65 | 0.068 | ° | 2.53 | 0.092 | ° |
| Playford High - Playford Low | -2.36 | 0.145 | ns | 2.67 | 0.068 | ° | 2.79 | 0.058 | ° |
| Bournemouth Low - Haikou Low | -3.40 | 0.007 | ** | 3.54 | 0.005 | ** | 4.10 | 0.001 | *** |
| Bournemouth Low - Playford Low | -1.74 | 0.405 | ns | 3.53 | 0.005 | ** | 4.30 | 0.000 | *** |
| Haikou Low - Playford Low | 1.59 | 0.447 | ns | 0.14 | 1.000 | ns | 0.39 | 0.790 | ns |

The difference in diversity between ‘low’ and ‘high’ complexity vegetation soils in Playford compared to Bournemouth and Haikou may be due to Playford’s relatively drier climate and the tendencies of native vegetation in this part of Australia to prefer relatively arid conditions. Such conditions are less conducive to supporting high microbial biodiversity (Delgado◻Baquerizo et al. 2018). In these drier environments, areas of lower vegetation complexity, such as urban lawns, are often heavily watered and fertilized. This practice can lead to higher nutrient loads relative to higher vegetation complexity native soils, potentially increasing microbial diversity independent of vegetation complexity. However, we note that our previous work with the Playford samples (Mills et al. 2020) indicated a consistent pattern in alpha diversity as found here in Bournemouth and Haikou (i.e. more vegetation complexity associated with greater bacterial alpha diversity). Although, our earlier study reported data from the V1-3 region of the 16S rRNA gene, rather than the V4 region reported here. As such, future work should further explore the effect of marker choice on vegetation-bacterial diversity associations.

### Differences in bacterial composition between vegetation complexities vary between cities

We next tested relationships between soil bacterial composition at the genus-level and the vegetation complexity of urban green spaces. The composition of bacterial communities was significantly different between ‘low’ and ‘high’ complexity vegetation in both Haikou (Bray-Curtis, F = 4.05, Pr(>F) < 0.05; Jaccard, F = 3.19, Pr(>F) < 0.01) and Playford (Bray-Curtis, F = 4.42, Pr(>F) < 0.05; Jaccard, F = 3.22, Pr(>F) < 0.05) (Table 2 & Figure 2b). However, Bournemouth had no significant difference between the vegetation complexities for both Bray-Curtis and Jaccard distances (Figure 2b).

**Table 2.** Main and pairwise PERMANOVA on soil bacterial genus communities for vegetation complexity nested within cities. Significance codes Pr(>F): ‘ns’ not significant; ‘°’ P < 0.10; ‘*’ P < 0.05; ‘**’ P < 0.01; ‘***’ P < 0.001.

| Formula = distance ~ City/Vegetation complexity |  |  |  |  |  |  |  |
| --- | --- | --- | --- | --- | --- | --- | --- |
| Main PERMANOVA |  | Bray-Curtis |  |  | Jaccard |  |  |
|  |  | R<sup>2</sup> | F | Pr(>F) | R<sup>2</sup> | F | Pr(>F) |
| City | df<sub>2,26</sub> | 0.52 | 19.61 | *** | 0.39 | 10.68 | *** |
| City/Vegetation complexity | df<sub>3,26</sub> | 0.13 | 3.25 | *** | 0.14 | 2.56 | *** |
| Pairwise PERMANOVA |  | Bray-Curtis |  |  | Jaccard |  |  |
|  |  | R<sup>2</sup> | F | Pr(>F) | R<sup>2</sup> | F | Pr(>F) |
| Bournemouth High - Haikou High | df<sub>1,9</sub> | 0.30 | 3.82 | ** | 0.23 | 2.70 | * |
| Bournemouth High - Playford High | df<sub>1,9</sub> | 0.60 | 13.36 | ** | 0.45 | 7.35 | ** |
| Bournemouth High - Bournemouth Low | df<sub>1,9</sub> | 0.16 | 1.78 | ns | 0.14 | 1.49 | ns |
| Bournemouth High - Haikou Low | df<sub>1,10</sub> | 0.42 | 7.19 | *** | 0.32 | 4.75 | ** |
| Bournemouth High - Playford Low | df<sub>1,9</sub> | 0.54 | 10.54 | ** | 0.40 | 5.95 | ** |
| Haikou High - Playford High | df<sub>1,8</sub> | 0.69 | 17.99 | ** | 0.53 | 8.94 | ** |
| Haikou High - Bournemouth Low | df<sub>1,8</sub> | 0.56 | 10.30 | ** | 0.43 | 6.13 | * |
| Haikou High - Haikou Low | df<sub>1,9</sub> | 0.31 | 4.05 | * | 0.26 | 3.19 | ** |
| Haikou High - Playford Low | df<sub>1,8</sub> | 0.61 | 12.58 | ** | 0.46 | 6.76 | ** |
| Playford High - Bournemouth Low | df<sub>1,8</sub> | 0.72 | 20.44 | ** | 0.56 | 10.35 | * |
| Playford High - Haikou Low | df<sub>1,9</sub> | 0.58 | 11.81 | ** | 0.45 | 7.34 | ** |
| Playford High - Playford Low | df<sub>1,8</sub> | 0.36 | 4.42 | * | 0.29 | 3.22 | * |
| Bournemouth Low - Haikou Low | df<sub>1,9</sub> | 0.61 | 13.86 | ** | 0.48 | 8.25 | ** |
| Bournemouth Low - Playford Low | df<sub>1,8</sub> | 0.67 | 16.22 | * | 0.52 | 8.52 | * |
| Haikou Low - Playford Low | df<sub>1,9</sub> | 0.49 | 8.74 | ** | 0.39 | 5.71 | ** |

Vegetation type (e.g., lawn, remnant woodland) is a known driver of microbial diversity and composition in urban soil (Hui et al. 2017; Mills et al. 2020). However, there is little consistency between soil microbial communities in what seem to be broadly similar ecological settings, as in our study, due to a complexity of multiple driving factors. Such factors include plant species turnover and soil properties that vary on broad spatial scales, such as temperature (Thompson et al. 2017) and, at finer scales, pH and salinity (Fierer & Jackson 2006). Certainly, pH and salinity have previously been found to strongly associate with urban soil bacterial community composition (Joyner et al. 2019; Mills et al. 2020). While we did not measure soil physicochemical properties here, they may, in some instances, override any effect of the vegetation community on the soil community and potentially lead to results as we saw in Bournemouth.

### Rare genera contribute to differences in community structure

We performed differential abundance testing to investigate which genera may have been driving the differences between the ‘low’ and ‘high’ complexity vegetation soils. Rare genera (i.e., < 1 % relative abundance) dominated the significantly differentially abundant bacteria between ‘low’ and ‘high’ complexity vegetation soils. For example, in Bournemouth, *Bacillus* (characteristic of ‘high’ complexity vegetation soils) was the only genus out of seven differentially abundant genera (P < 0.05, Figure 3) that was also dominant in relative abundance (> 1 % relative abundance, Figure 2c) between the vegetation complexity levels – the other six genera were less than 1 % in relative abundance. In Haikou, *Rubrobacter* (characteristic of ‘low’ complexity vegetation soils) was the only genus out of four differentially abundant genera to also be greater than 1 % in relative abundance (P < 0.05, Figure 3), and in Playford, *Flavisolibacter* and *Gemmata* (both characteristic of ‘low’ complexity vegetation soils) were the only differentially abundant genera out of twenty-three that were also dominant (P < 0.05, Figure 3). Overall, differential abundance tests showed that there are soil bacteria characteristic of either ‘low’ or ‘high’ complexity vegetation within each city and that rare taxa are important in defining these communities. This result is consistent with other findings that rare bacteria biogeographically distinguish forensic soil samples (Damaso et al. 2018; Habtom et al. 2019).

**Figure 3.**
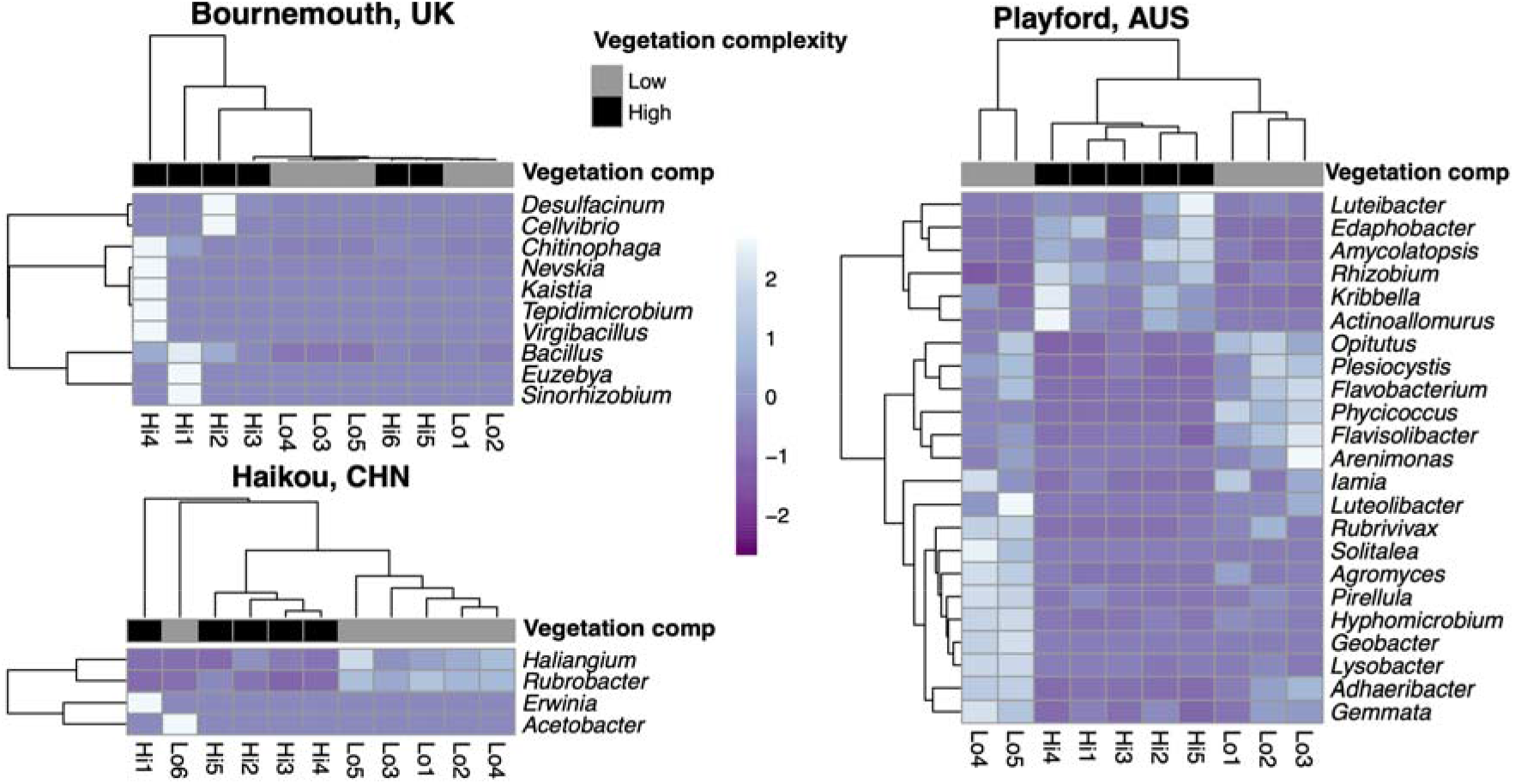
Differentially abundant bacterial genera from Bournemouth, Haikou, and Playford measured by log-2 fold-change with P-value < 0.05. Extreme ends of the heat color scale represent 3 standard deviations from the mean rarefied abundance for each genus across samples. Hierarchical clustering of genera (rows) and vegetation replicates (columns) are both by Manhattan distance.

The rare genera (< 1 % relative abundance) that dominated the differential abundance testing between the vegetation complexities (P < 0.05, Figure 3) were potentially functionally important to these locations as has been found in greenhouse soils (Xue et al. 2020). For example, *Luteibacter* was the only genus to significantly represent ‘high’ vegetation complexity across all cities in this study and species of this genus are known to live both in soil and on humans (Kämpfer et al. 2009; Akter & Huq 2018). The ‘high’ complexity vegetation soils of Bournemouth had significantly more *Sinorhizobium* (nitrogen-fixers; Mitsui et al. 2004) and *Kaistia* (methanotrophs; Im et al. 2004) of the order Rhizobiales (Garrido-Oter et al. 2018) than ‘low’ complexity soils. However, most differentially abundant genera in Bournemouth were higher in only one site relative to others; therefore, they are not characteristic of either vegetation complexity studied here. In Haikou, *Haliangium* (producer of fungicidal haliangicins; Fudou et al. 2001), *Rubrobacter*, and *Acetobacter* were significantly more abundant in the ‘low’ than in the ‘high’ vegetation diversity soils, whereas *Erwinia* (genus of many plant pathogen species; Barras et al. 1994; Vanneste 2000) was more abundant in the ‘high’ vegetation diversity soils. In Playford, *Rhizobium* (nitrogen-fixers; Garrido-Oter et al. 2018) were significantly more abundant in the ‘high’ complexity vegetation.

The differential abundance findings that imply rare genera are driving the community differences are further supported by the similarity between Bray-Curtis and Jaccard ordinations (Figure 2b). Further, ordinations of only the rare genera (those < 1 % relative abundance) were similar to ordinations using the whole community (Figure S3), therefore implying that rare genera are driving these patterns; however, these data were over-dispersed. Additionally, the rank-abundance curve showed there were 300 of 318 genera with less than 1 % relative abundance across all sites (Figure S1), and, when pooled, had an average relative abundance of 35 % across all sites (Figure 2c). These findings indicate that rare genera may be quite valuable to urban soils and that they shouldn’t be overlooked when planning soil microbial conservation. To that end, rare microorganisms have been identified to play key functional roles, from biogeochemical cycles to holobiont health (Jousset et al. 2017). Further exploration of the functional contribution of rare bacteria in urban green spaces would provide deeper understanding of their value in conservation and restoration efforts.

### Conclusions

Our study suggests that a global comparison of cities in terms of vegetation factors driving microbial diversity may be limited due to the overall strength of their differences driven by geographic or climatic factors. However, investigating trends related to vegetation complexity within cities may produce general recommendations about fostering microbial biodiversity. Certainly, rare taxa should not be overlooked when considering the conservation of microbial biodiversity. Urban green space design for conservation of microbial biodiversity, biogeochemical cycling, public health outcomes and public usability will likely require complementary proportions of both ‘low’ and ‘high’ complexity vegetation green spaces. However, what those proportions are will need to be investigated on a city-by-city, or region-by-region basis.

Restoration of biodiversity in urban green spaces has the potential to build native microbial communities. Such endeavors will require local adaptive management within urban green space landscapes that will allow practitioners to understand the knowledge gaps pertaining to their city and properly investigate the outcomes of their efforts (Gellie et al. 2018). Local knowledge gaps may include: understanding functional microbial biodiversity; examining how differently designed green spaces influence environmental and human microbiota (e.g., ‘low’ and ‘high’ complexity, native and novel species mixtures); and determining if the use of remnant inoculations accelerates the recovery of native microbial phylogenetic and functional diversity. Further, there is currently a strong call to ‘decolonize’ public spaces in colonial and imperial countries (Parker 2018; Giblin et al. 2019). Therefore, it would be interesting to track whether such cultural modifications to urban designs influences environmental and human microbiota given that ‘low’ and ‘high’ complexity green spaces are somewhat representative of these cultural differences. More work is needed to describe the functional contributions of rare bacteria in urban soils and to determine the best ways to conserve and restore microbial biodiversity to provide the breadth of ecosystem services that they could provide to the urban landscape.

## Author statements

### Author contributions

JM, CAS, LW, PW, AL, MB: conceptualization. JM, CAS, CS, LW, JY, EM, SY, VY, MB: investigation. JM, CAS: data curation. JM, CAS, LW, JY, TT: formal analysis. PW, MB: funding acquisition. JM, CAS, LW, JY, TT, AL, MB: methodology. JM, CAS: software. JM, CAS, LW, CS, PW, AL, MB: project administration. JM, LW, EM, CS, SY, VY, AL: resources. LW, CS, TT, PW, AL, MB: supervision. All: validation. JM: visualization. JM: writing – original draft. All: writing – review & editing.

## Acknowledgments

Michael Rowland (Bournemouth, Christchurch and Poole Council) and Soumya Prasad (Jawaharlal Nehru University, New Delhi) for assistance in selecting sites in Bournemouth and New Delhi (even though this dataset was not used), respectively. Jo Park, Ken Daniel and the City of Playford for assistance with site selection and access. Qiyong Liu and Keke Liu from the Chinese CDC, and Peng Bi (University of Adelaide) for coordinating and providing samples from China.

