## Supplementary Material for "Rare genera differentiate urban green space soil bacterial communities in three cities across the world"

^D^ Deceased

**Table S1** Bacterial 16S rRNA V4 region forward and reverse primers.

| Primer | Primer sequence | Citations |
| --- | --- | --- |
| 515F | AATGATACGGCGACCACCGAGATCTACAC TATGGTAATT GT GTGCCAGCMGCCGCGGTAA | Caporaso et al. (2011); Caporaso et al. (2012) |
| 806R X-barcode | CAAGCAGAAGACGGCATACGAGAT XXXXXXXXXXXX AGTCAGTCAG CC GGACTACHVGGGTWTCTAAT |  |

**Table S2** Custom sequencing primers.

| Read 1 sequencing primer | Primer sequence |
| --- | --- |
| Forward primer pad | TATGGTAATT |
| Forward primer linker | GT |
| Forward primer | GTGYCAGCMGCCGCGGTAA |
| Read 2 sequencing primer | Primer sequence |
| Reverse primer pad | AGTCAGCCAG |
| Reverse primer linker | CC |
| Reverse primer | GGACTACNVGGGTWTCTAAT |
| Index sequencing primer | AATGATACGGCGACCACCGAGATCTACACGCT |

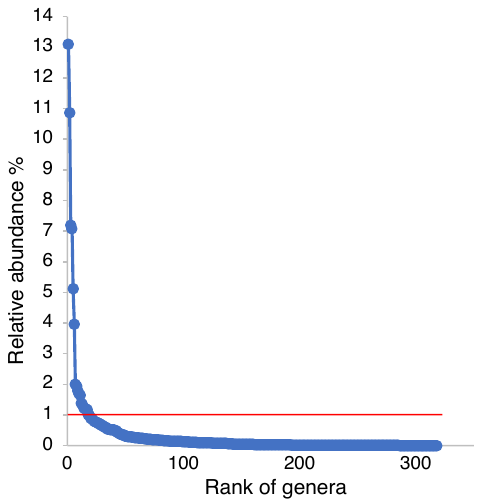

**Figure S1** Rank-abundance curve of bacterial genera to determine rare genera cut-off. Red line shows the curve’s approximate inflection at 1 % relative abundance.

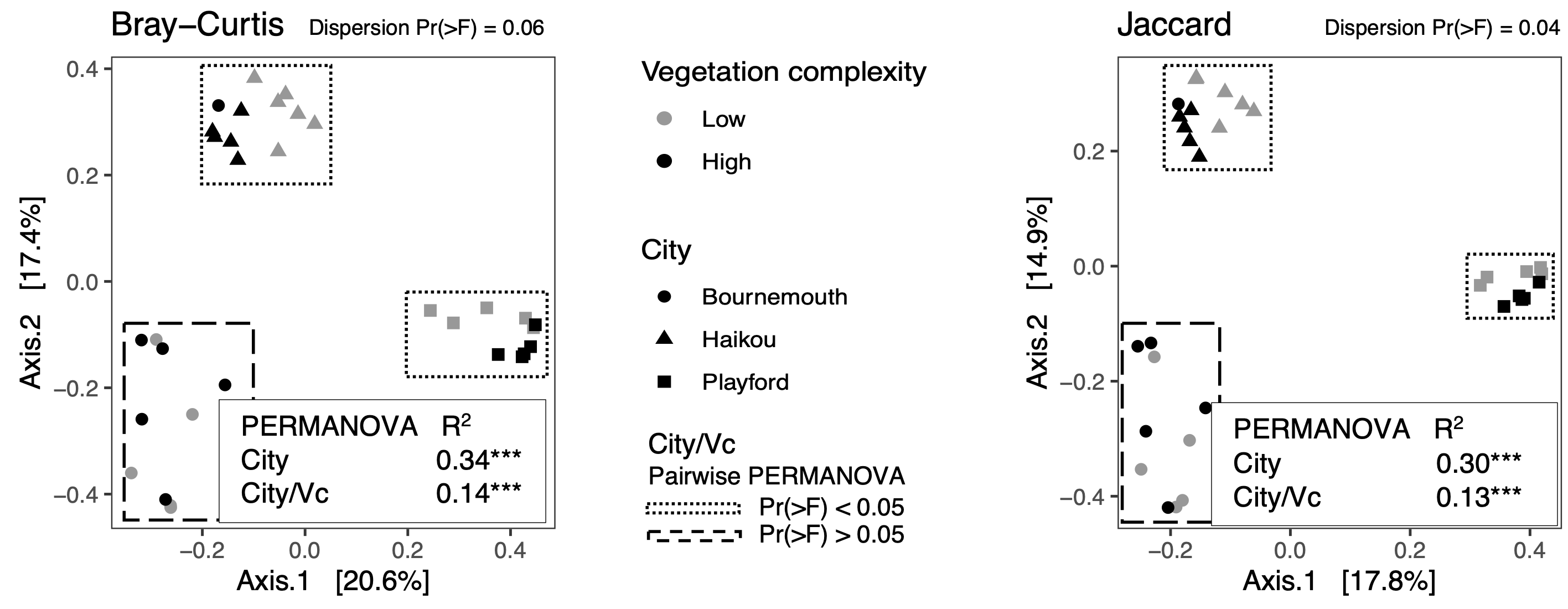

**Figure S2** ASV-level ordinations. Jaccard is over-dispersed and Bray-Curtis is very close to it.

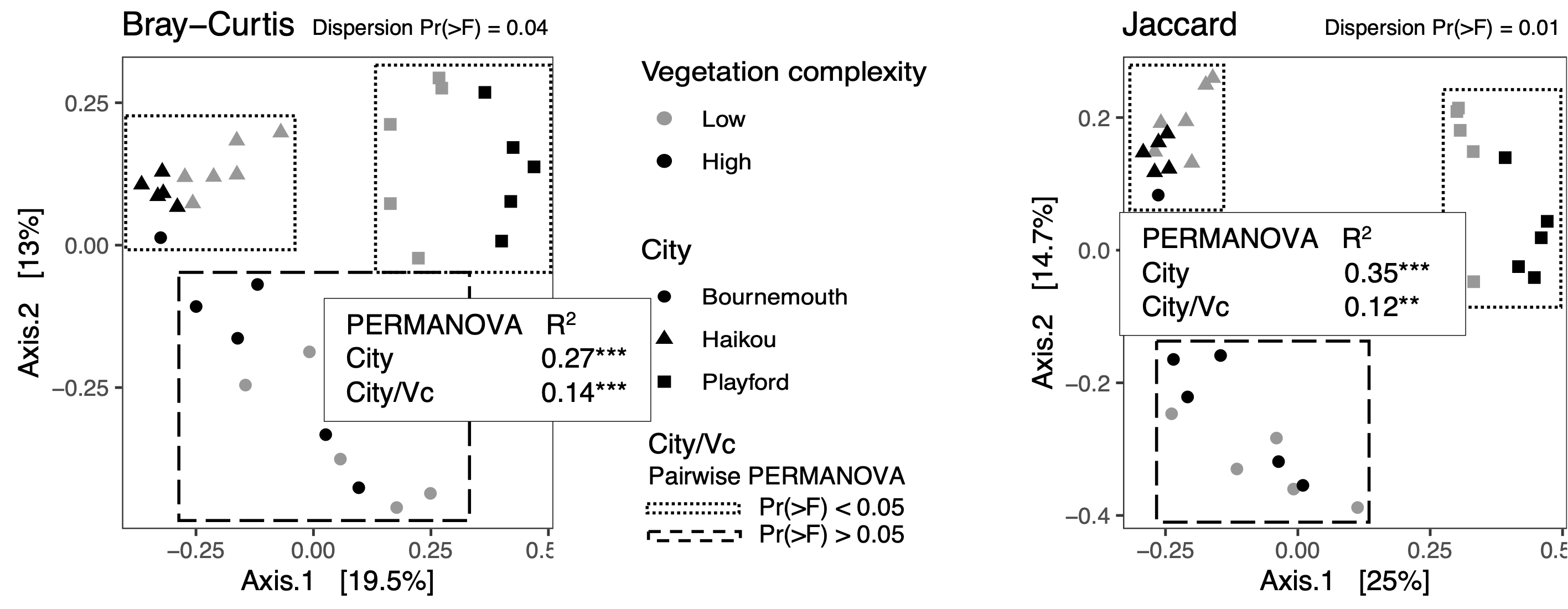

**Figure S3** Ordinations of rare genera only (those < 1 % relative abundance). Both datasets are over-dispersed.
